# PCL-seq: enhanced high-resolution transcriptomic profiling of region of interest in fresh frozen and FFPE tissues

**DOI:** 10.1101/2024.08.05.606746

**Authors:** Xue Dong, Xiaonan Cui, Mengzhu Hu, Wenjian Zhou, Weiyang Shi

## Abstract

The spatial heterogeneity of gene expression has propelled the development of multiple spatial transcriptomics technologies. Here, we present **p**hoto**c**leavage and **l**igation sequencing (PCL-seq), an method for spatial indexing using a light-controlled DNA labeling strategy on tissue section. PCL-seq uses photocleavable oligonucleotides and ligation adapters to construct transcription profiles of region of interest (ROI), selected by microscopically controlled photo illumination apparatus in tissue sections. Applied to mouse embryos, PCL-seq obtains gene expression matrices that align with spatial locations and competitive data quality, featuring around 1.7×10^5^ UMIs and 8,600 genes (irradiation diameter=100µm). PCL-seq can also apply to formalin fixation and paraffin embedding (FFPE) mouse embryo sections, whereas obtained competitive data output and recovered thousands of differentially enriched transcripts from limb and skeleton. Additionally, PCL-seq can achieve subcellular resolution, which was demonstrated for differential expression between nuclear and cytoplasmic. Thus, PCL-seq provides an accessible workflow for spatial transcriptomic analysis in frozen and FFPE tissue at subcellular resolution.

## Introduction

With the advancement of single cell sequencing technologies, it has been recognized that many gene, in biological systems, need to be properly regulated in space for the system to function^1–6^. Thus, the analysis of spatial environment is crucial for inferring deeper biological significance. Spatial transcriptome technology combined with conventional single cell sequencing technology, in situ technology and other omics technologies can enable the study of gene expression in spatial location to be combined to the ultra-high-resolution single cell level, which has promising applications in the fields of cancer, immunity, neurology, and development^7–11^.

Existing spatial transcriptome technologies can be divided into two categories. One approach involves performing unbiased transcriptome analysis across the entire tissue, mainly including methods based on in situ hybridization^12–15^ and in situ capture^5,16–20^. Another strategy in spatial transcriptomics research is to selectively isolate and analyze regions of interest (ROIs) with known locations and morphologies.^21^. In practical applications, compared to whole transcriptome analysis, the analysis of only specific regions is easier to focus on the target and is more suitable for clinical sample studies of tissue heterogeneity^22^.

ROIs acquisition can be performed by physical or optical marking. Physical isolation methods include LCM-seq^23^ or Geo-seq^24^, which cut small group of cells or a single cell in a tissue section with laser, followed by RNA-seq. However, the use of precision instruments limits the versatility and resolution of such techniques. In contrast, the strategy of separating ROIs based on optical marking is not constrained by the use of sophisticated instruments like laser scanning confocal microscope. This approach typically involves labeling tissues with photosensitive compounds that undergo molecular changes in response to light exposure, thereby distinguishing illuminated from unilluminated areas for subsequent analysis. Commercial platforms such as GeoMx Digital Spatial Profiling^25^ (DSP) use UV linker as photosensitive compounds. Upon light exposure, these linker breaks and releases oligos containing protein and mRNA abundance information into the solution. These oligos are then collected by micro-pipettes and sequenced to determine the expression levels of each target. Currently, DSP can simultaneously analyze spatial information of over 180 proteins and 18,000 genes. The limitation of DSP lies in its use of probes to human or mouse transcriptome, which restricts its application in other species. Other techniques such as ZipSeq^26^, PIC^27^, and Light-seq^28^ also employ different photosensitive compounds to perform transcriptome analysis of ROIs. However, the 6-nitropiperonyloxymethyl (NPOM) compounds used in ZipSeq and PIC are costly, and the 3-cyanovinylcarbazole nucleoside (CNVK) utilized in Light-seq has not yet been commercialized, thus limiting the general applicability of these techniques.

Here, we introduce a method termed Photocleavage and Ligation Sequencing (PCL-seq), which employs photocleavable (PC) linker compounds as photosensitive groups to obtain gene expression profiles of arbitrary ROIs in tissue sections. This approach offers a more economical and versatile solution for transcriptome analysis. We demonstrate that PCL-seq not only obtains high-depth spatial transcriptome profiles from conventional fresh frozen sections, but also work on FFPE sections of mouse embryo. In addition, PCL-seq can achieve subcellular resolution to profile gene expression in subcellular structure.

## Results

### PCL-seq overview

PCL-seq utilizes a light-controlled spatial transcriptome labeling technique for targeted transcriptome analysis in specific ROI within tissue sections. This method involves the use of reverse transcription (RT) primer modified with a PC linker to first convert mRNA within the tissue section to cDNA with in situ reverse transcription. Subsequently, patterned UV light cleaves the PC linker, exposing the 5’ end of cDNA with a phosphate group.^25,29^. Then, an adaptor is attached to the cDNA within the chosen ROI through a ligation reaction. In areas where there is no UV irradiation, the unbroken PC linker will effectively hinder cDNA ligation to the adapter (Figure 1). This enables precise localization and analysis of the transcriptome in designated tissue areas.

**Figure 1.**
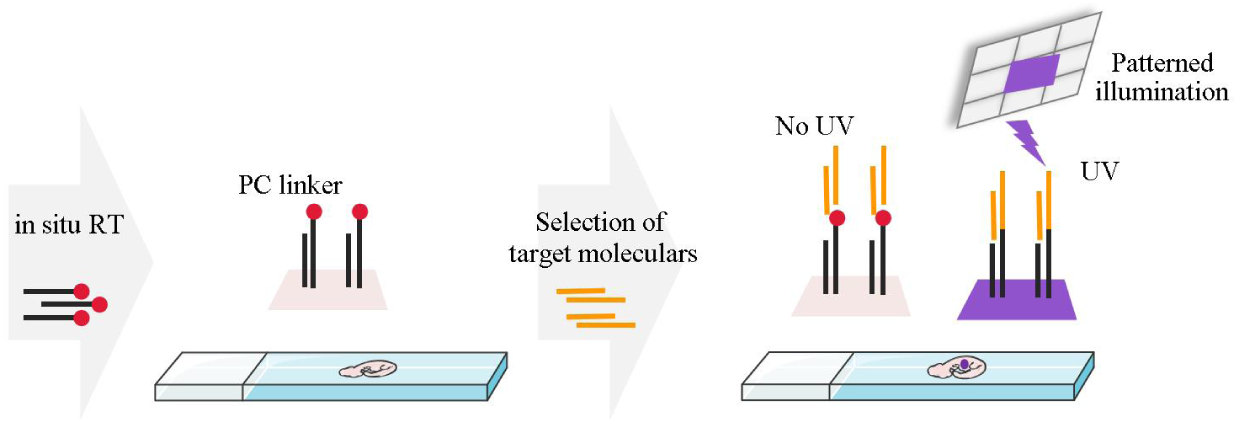
Schematic for light-controlled DNA attachment strategy. Frozen tissue sections adhered to poly-L-lysine slides were fixed and permeabilized, followed by an in situ RT reaction utilizing PC linker-modified RT primers to synthesize ‘locked’ cDNA. UV light, selectively filtered by Mosaic’s digital mirror device system, was applied to the tissue sections to illuminate regions corresponding to the ROI. In the illuminated regions, the PC linkers on the newly synthesized cDNA were cleaved, allowing the ligation of sequencing adapters. Consequently, cDNA in these regions could be specifically amplified using primers complementary to the adapters, thus enabling the construction of ROI-specific transcription libraries. In contrast, the PC linkers in the non-illuminated regions remained intact, thereby preventing the ligation of adapters to the cDNA and subsequently inhibiting the amplification of cDNA from these areas.

The key to PCL-seq is the specific amplification of cDNA from the illuminated region. To achieve this objective, we have implemented the following optimizations. In PCL-seq, the Andor Mosaic system, which is a digital mirror devices (DMD) based system that can illuminate computer generated patterns on sides with a resolution of 2 um ( when using 20x objective), is used to perform the function of selecting ROIs. Therefore, we first need to ensure that the UV light filtered through the mosaic is reflected as UV beams conforming to the shape of each ROI, precisely illuminating the respective ROI area. We optimized the intensity and duration using photo-dissection experiments on functionalized slides. This involved using amino-modified RT-PC linker-FAM primers covalently immobilized on the surface of CodeLink Activated slides^16,30^. These slides were subjected to various light exposure durations (10, 20, 30 seconds) and intensities (10%, 40%, 70%, 100% power) to determine the optimal settings for specific area illumination (Supplementary Figure 1A). The key criterion for selecting the optimal conditions was the complete disappearance of the green fluorescence signal, indicating effective PC linker detachment, which was most pronounced at 100% light intensity and 10 seconds of illumination time. These parameters ensured that the cleavage of the PC linker occurred exclusively in the irradiated areas, thereby enhancing the specificity and reducing contamination from neighboring areas.

Secondly, during the in situ tissue reactions phase of PCL-seq, it is critical to highlight that the presence of the PC linker on the RT primers does not inhibit the hybridization between complementary strands. This characteristic allows sequences in non-irradiated areas to hybridize with sequencing adapters. Consequently, these hybridized sequences can act as “bridges” inadvertently facilitating the amplification of transcript information from non-targeted areas during the subsequent amplification process. To address the aforementioned impact, we use a high concentration of formamide to wash the cells or tissue sections after the ligation reaction, thereby eliminating hybridization products between the ligation adaptors and cDNA in the non-irradiated regions, thus reducing background noise. We utilized functionalized slides and fluorescent oligonucleotides to simulate the in situ reaction process, incorporating formamide washing to visualize key molecular events at each step. This approach allowed us to directly assess the feasibility of specifically labeling ROI regions. Specifically, CodeLink slides modified with RT-PC linker-FAM oligonucleotides were subjected to sequential illumination in designated areas, adapter ligation, and high-concentration formamide washes. Microscopy was used to record the changes in fluorescence signals at each step of the reaction (Supplementary Figure 1B). The fluorescence results demonstrated the stability of optimized illumination conditions and underscored the necessity of formamide usage. Overall, these results indicate that we have preliminarily achieved the precise and specific fluorescent labeling of molecules in photo illuminated regions, which is a crucial prerequisite for the feasibility of the PCL-seq.

### Establishment of PCL-seq for ROI-specific expression profiling

Following these optimizations, we established PCL-seq for the analysis of the spatial transcriptome in ROI. Experimentally, we began by using PC linker-modified RT primers to perform an in situ RT reaction on fixed and permeabilized tissue sections, synthesizing first-strand cDNA. We then utilized the Mosaic companion software to visualize and select the ROI, followed by specific UV light irradiation of the designated area. The PC linker responded to the UV irradiation by detaching from the newly synthesized cDNA, allowing for the ligation of sequencing adapters. The formation of hybrids in the non-irradiated regions made high concentration formamide washing particularly critical. Post-washing, the stable ligation products in the irradiated areas were retained. Finally, after enzymatic digestion of the sample, the cDNA was isolated, amplified, and library prepared for next-generation sequencing (Figure 2A and Supplementary Figure 2).

**Figure 2.**
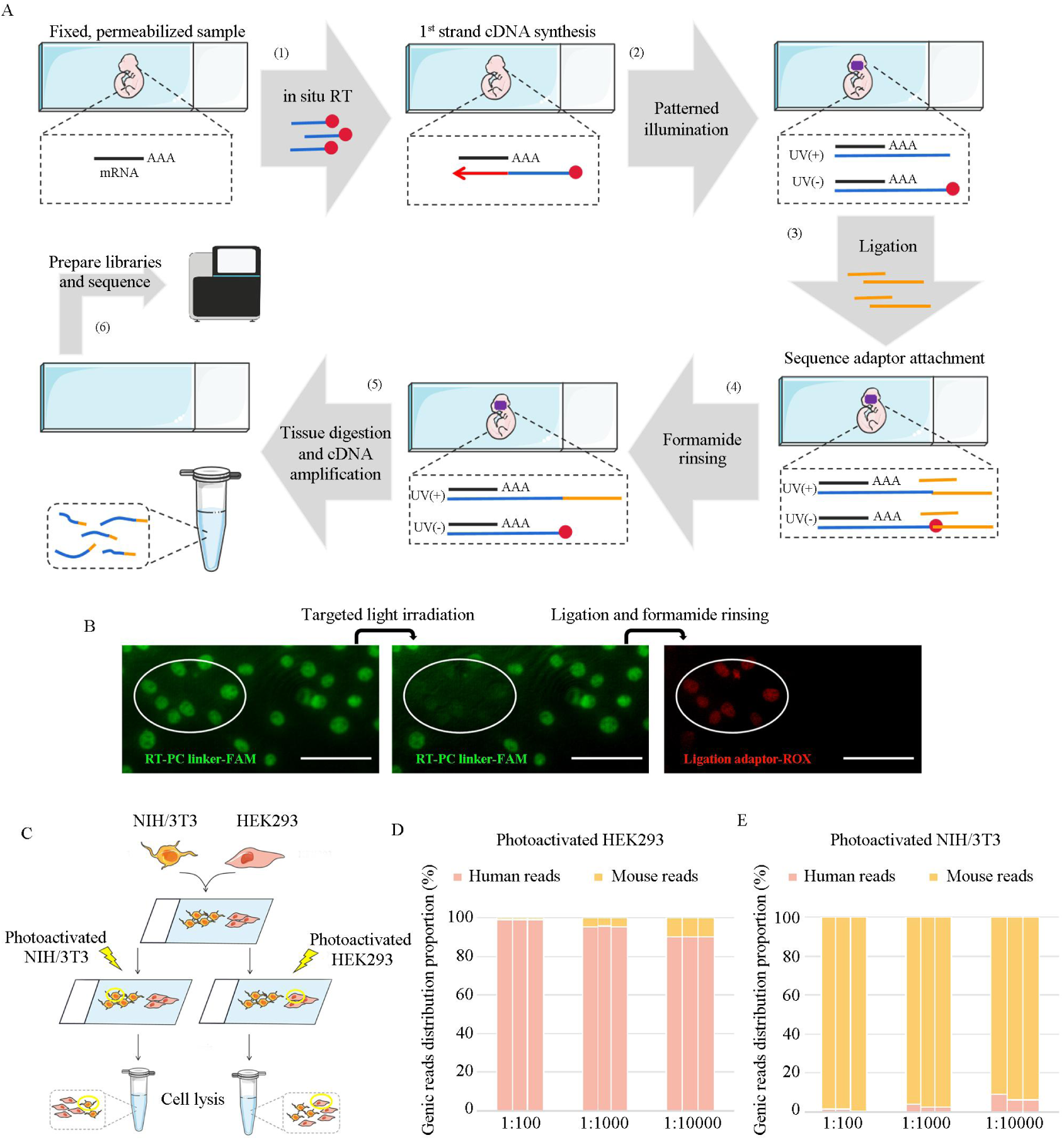
Establishment of PCL-seq for ROI expression analysis. (A) PCL-seq workflow: (1) First-strand cDNA was synthesized by in situ RT of fixed and permeabilized tissue sections using RT-PC linker primers. (2) The ROI was subjected to patterned UV irradiation. (3-4) Sequencing adapters were introduced into the cDNA molecule within the ROI via a ligation reaction. (5) The sample undergoes enzymatic digestion and purification to isolate the adaptor attached cDNA fragments. (6) The sequencing library was generated after amplification and fragmentation of the cDNA molecules. (B) The NIH/3T3 cell line was stained with RT-PC linker-FAM primers through an in situ RT reaction and imaged following photo-cleavage, ligation, and formamide rinsing. The elliptical area denotes the photo-activated region. Scale bar, 100 μm. (C) Testing the precision of PCL-seq by specifically photo-tagging co-cultured HEK293 and NIH/3T3 cells. (D, E) Background levels from non-irradiated cells were assessed using human–mouse mixed cultures. N=3 replicates.

Next, we validated PCL-seq’s ability to specifically acquire ROI transcript information on cell cultures. First, we confirmed the specific labeling of cells in the illuminated area using a fluorescently labeled barcode adapter. NIH/3T3 cells were fixed, permeabilized, and subjected to in situ RT reactions with RT-PC linker-FAM primers. Subsequently, we illuminated a small region of the cells using the mosaic system and added a red fluorescently labeled adaptor. Indeed, this approach effectively achieved the specific labeling of cDNA molecules in the illuminated region on cell cultures (Figure 2B). Meanwhile, having established specific attachment of adaptor to cDNA in photo-illumianted areas, we set to test specificity and sensitivity for PCL-seq. We first performed the following species mixing experiment with cultured human and mouse cell lines. We selectively illuminate a small portion of one cell species in the background of the other species which are un-illuminates, at different ratio. To do this, we co-cultured HEK293 cells (Human) and NIH/3T3 cells (Mouse) on distinct regions on the same slide (Figure 2C). Upon fixation and in situ reverse transcription, selective photo-illumination was applied to distinct zones within the co-culture, resulting in three experimental conditions: 1) photo-illuminated HEK293 area; 2) photo-illuminated NIH/3T3 area; 3) non-photocleaved (control). The number of cells illuminated compared to the un-illuminate species is of 1/100, 1/1000, and 1/10000. Sequencing reads were mapped to a combined human and mouse reference genome to ascertain the proportion of genic reads from each species under different experimental conditions. The libraries derived from samples without UV irradiation (condition 3) gave negligible read counts (control reads vs experimental reads=4.8 ×10^3^ and 5.1 ×10^6^), suggesting unillumated cells fail to produce sequencing library reads. For HEK293 illuminate experiment, the proportion of human genic reads was 99.08%±0.07%, 95.38%±0.1%, and 90.06%±0.06%, with the remaining arising from mouse genome, for the scales of 1/100, 1/1000, and 1/10000 of the irradiated area to the total area, respectively (Mean±SD, Figure 2D). In converse experiment whereas NIH/3T3 cells are illuminated, mouse genic reads constituted 98.96%±0.56%, 97.01%±0.76%, and 92.84%±1.7% of the total reads for the same ratio (Mean±SD, Figure 2E). These results demonstrate that our PCL-seq method effectively captures mRNA from targeted cells, even when as few as 0.01% of the cells in the total population are selected.

### PCL-seq for fresh frozen tissue sections

We used mouse embryo to test PCL-seq for specific ROI expression analysis on tissue sections. First, PCL-seq demonstrated the ability to specifically label cDNA molecules in the target region even against a complex tissue background, consistent with its performance on cell cultures (Supplementary Figure 3A). Next, we tested PCL-seq’s ability to obtain gene expression profiles from cells at specific spatial locations. Specially, adjacent E13.5 mouse embryo sections were fixed with formaldehyde and underwent reverse transcription. Next, sections were subjected to immunostaining for *PAX6*, a major gene in eye development and widely expressed in the neuroectoderm and epidermal ectoderm during eye development^31,32^ to mark the eye region. Then, PAX6-positive and PAX6-negative domains were independently photo-irradiated (n=4, 4 replicates respectively, Figure 3A), using a ×20 objective lens (irradiation diameter: 100 μm). The sections were then lysed and cDNA libraries were amplified and sequenced. At an average of 4.5×10^6^ reads per sample, PCL-seq detected an average of more than 7,500 genes from above two regions, and UMI yield varied for different regions depending on cell and RNA content of each region, ranging from 1,000 to ∼1,600 UMIs per 10×10µm^2^ unit area (Figure 3B, C).

**Figure 3.**
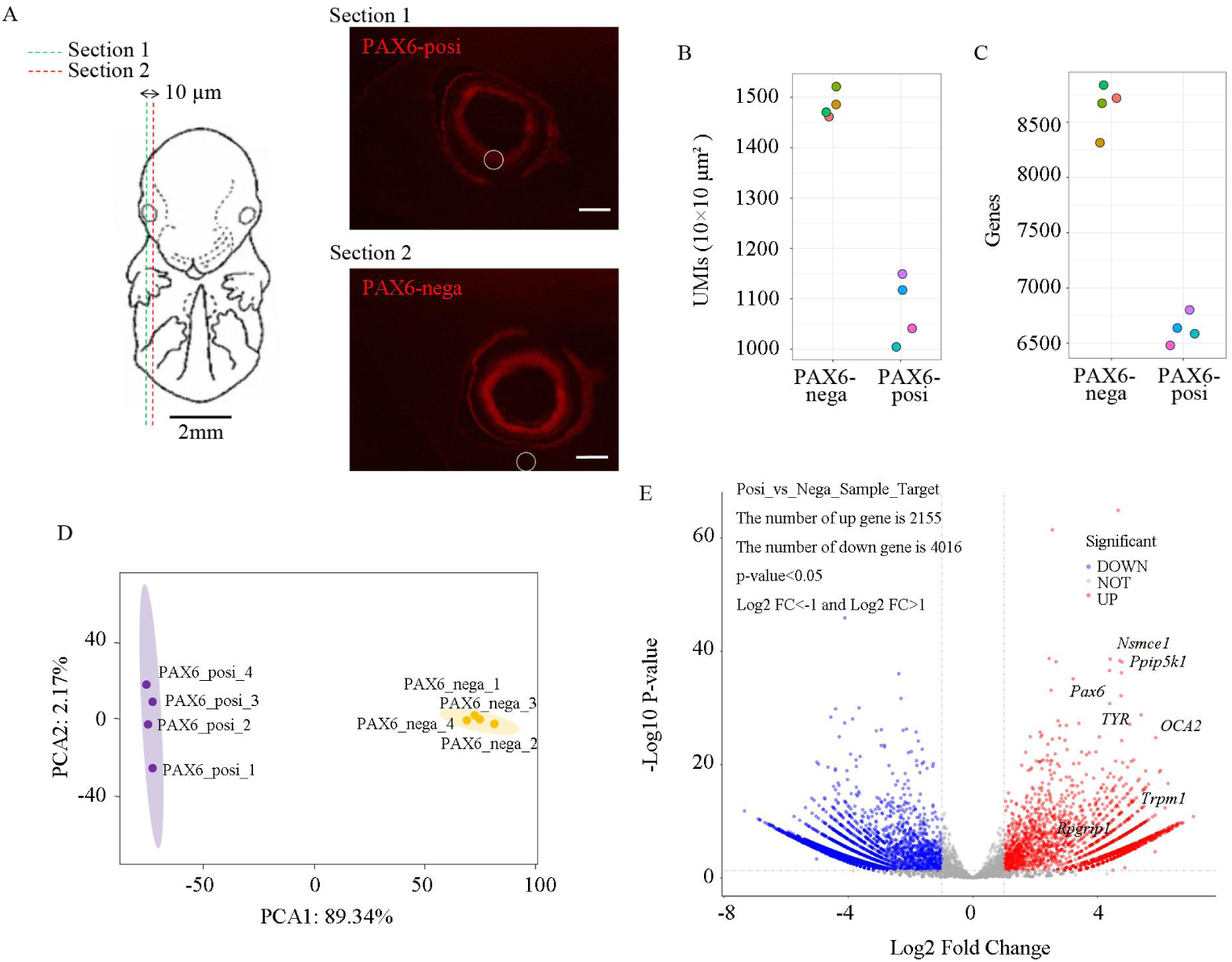
Analysis of PCL-seq for ROI-specific profiling in fresh frozen mouse embryo sections. (A) Left: Schematic illustration of adjacent sections in an E13.5 mouse embryo. Scale bar, 2mm. Right: Images of immunofluorescence staining using Alexa Fluor ○R 647 anti-PAX6 antibody (red). Mouse embryonic sections of PAX6-positive or -negative cells were photo-irradiated. The irradiated area is a circle with a diameter of 100 μm. Scale bar, 200 μm. (B, C) Numbers of detected per-area UMIs (B) and genes (C) are shown. (D) Two-dimensional PCA for the expression profiles is shown. (E) DEGs are clustered and shown in volcano plots.

We will further explore whether the gene expression information obtained through PCL-seq originates from the cells in the light-irradiated region. Four replicates of each area were strongly correlated (Pearson correlation coefficient >0.9, Supplementary Figure 3B). Firstly, we used DESeq2 (ref. ^33^) to perform the expression analysis of *GAPDH* and *PAX6* genes in PAX6-positive and PAX6-negative regions, respectively (Supplementary Figure 3C, D). The results demonstrated a resemblance to *PAX6* antibody staining, revealing that the *PAX6* gene expression was over twice as high in the positive region compared to the negative region. Conversely, *GAPDH* was detected in both the positive and negative regions, with no significant difference in its expression between the two regions. These results can provide preliminary evidence for the correlation between gene expression and spatial localization. Next, we conducted a comprehensive differential gene analysis between the two regions. Two-dimensional principle component analysis (PCA) indicated that the expression profiles were distinct (Figure 3D). And DEG analysis revealed 2,155 genes were up-regulated and 4,016 genes were down-regulated in PAX6-positive vs. PAX6-negative regions by heat maps (Supplementary Figure 3E) and volcano plots (Figure 3E). In addition to the *PAX6* gene, we also detected the up-regulation of genes such as *Pmel*, *Trpm1*, *TYR*, *Rpgrip1*, and *OCA2* in the PAX6-positive region. These genes are closely related to eye development and their functions have been previous documented^34–36^. Additionally, PCL-seq identified the expression of genes such as *Ppip5k1* and *Nsmce1* in the up-regulated gene matrix, whose role in mouse embryonic eye has not been studies but specific expression pattern is corroborated by immunohistochemical staining (ISH) results from the EMAGE Gene Expression

Database of resources provided by e-Mouse Atlas Project (https://www.emouseatlas.org/emap/home.html) (Supplementary Figure 4A). *Ppip5k1* plays a significant role in cellular processes such as phosphate and bioenergetic homeostasis, though its biological significance is relatively less understood^37^. *Nsmce1* has been associated with neurological diseases, although its role in the development of the mouse embryonic eye has not been previously described^38^. In summary, the above results indicate that PCL-seq can specifically obtain transcriptome maps of tissue section ROIs.

### PCL-seq for formalin fixation and paraffin embedding tissue sections

Formalin-fixing and paraffin embedding (FFPE) samples is a commonly used method of storing pathological tissues in clinical practice^39^. However, the utilization of formalin and the impact of prolonged storage resulted in varying degrees of RNA degradation and diminished mRNA accessibility in FFPE samples. These factors collectively complicate the analysis and study of the transcriptome in such specimens^40^. Recently, several approaches for unbiased, spatially resolved transcriptomics of FFPE samples was developed^41,42^. In fact, the strategy of such techniques lies in improving mRNA accessibility through heat-induced crosslink reversal, thereby enabling transcriptome analysis of FFPE tissue sections. Combining PCL-seq with the decrosslinking, we explored the spatial differences in gene expression in mouse embryo FFPE sections. Adjacent sagittal E15.5 mouse FFPE sections were subjected to crosslink reversal before PCL-seq and the embryonic limb and skeleton region were photo-irradiated (n=4, 4 replicates respectively), using a ×20 objective lens ( irradiation diameter: 100µm) in the same way (Figure 4A). With average of 7.1×10^6^ reads per sample, we detected an average of 4787 genes and 5119 genes from the limb and skeleton, respectively (Figure 4B). Meanwhile, the number of UMIs detected in limb and skeleton were 546 and 424 per 10×10µm^2^ unit area on average (Figure 4C). PCL-seq has higher sensitivity compared to other spatial transcriptome techniques on FFPE sections, in which 138 UMIs were detected in mouse brain FFPE sections using 10×Visum commercial chip^41^ for tissue areas of same size.

**Figure 4.**
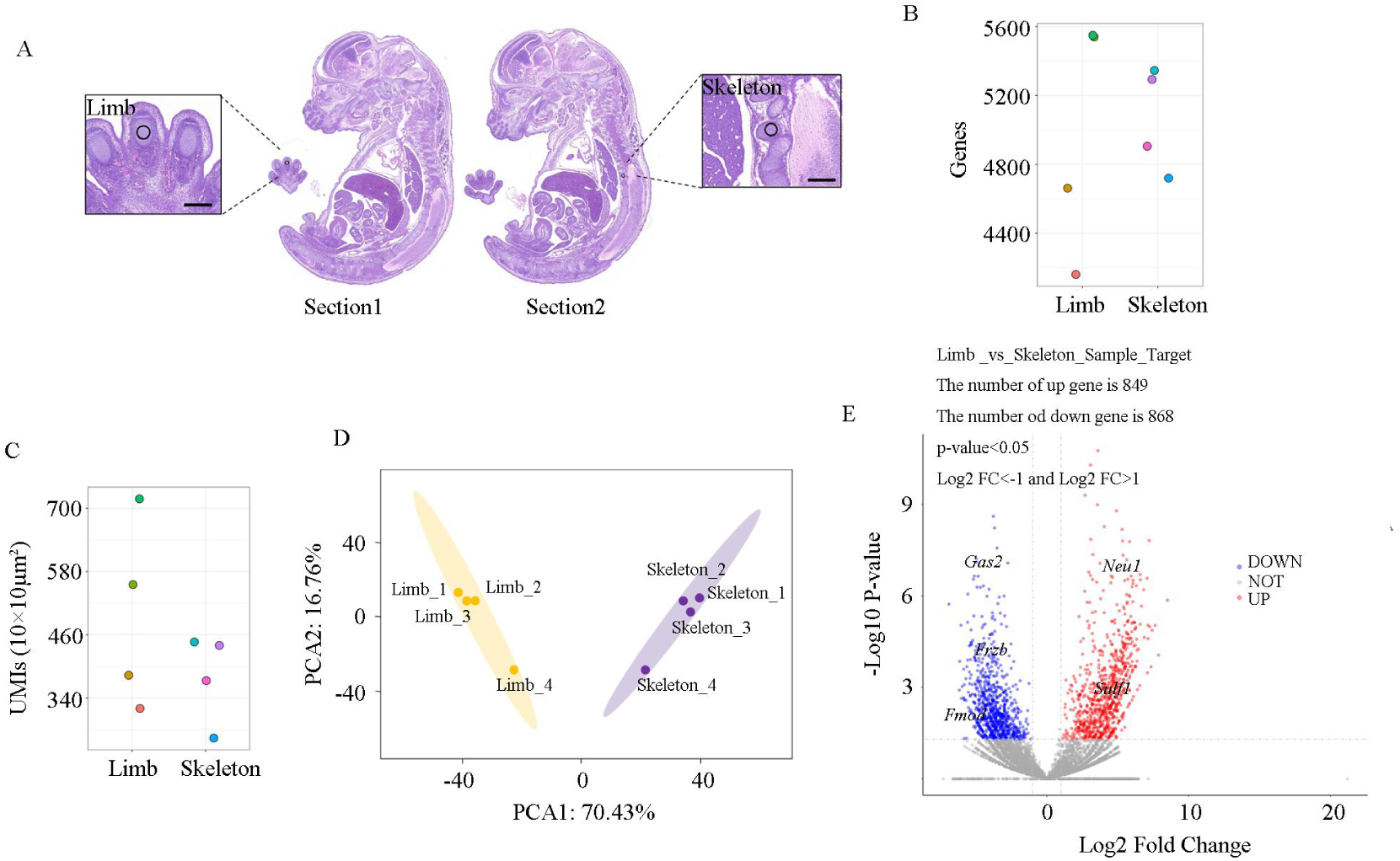
PCL-seq for formalin fixation and paraffin embedding mouse embryo sections. (A) Hematoxylin and eosin (H&E) staining images of adjacent mouse embryo sections. Limb and skeleton were separately photo-irradiated. The irradiated area is a circle with a diameter of 100 μm. Scale bar, 250 μm. (B, C) The number of detected genes (B) and UMIs per 10×10µm^2^ unit area (C) are shown. (D) Two-dimensional PCA for the expression profiles is shown. (F) DEGs are clustered and shown in volcano plots.

Next, PCA analysis showed that the expression patterns of limb and skeleton were clearly distinguished (Figure 4D). DEG analysis revealed a total of 1,717 genes between limb and skeleton (Figure 4E). Among them, *Sulf1* and *Fmod* was detected as limb or skeleton specific DEGs, and their spatial localization was also confirmed by stereo-seq^5^ study for mouse embryo. In addition, specific positional expression of *Neu1* detected in the limb, *Frzb*, which was detected in skeleton, was confirmed by ISH analysis from EMAGE Gene Expression Database as previously described (Supplementary Figure 4B). In summary, these results demonstrate that PCL-seq is also capable of obtaining transcriptomic profiles for specific ROI in FFPE samples. This indicates that it has broad applicability and significant utility in various research contexts.

### PCL-seq achieves subcellular resolution expression profiling

Subcellular resolution allows for precise localization of gene expression within different subcellular structure, which is crucial for understanding gene functions and regulatory mechanisms^43,44^. The resolution of PCL-seq is dependent on the size of the micro-mirrors in the DMD from Mosaic system, with a theoretical maximum resolution of up to 2 μm under a 20x objective lens. Therefore, we further investigated the potential of PCL-seq to obtain transcriptomic maps of subcellular structures. HeLa cells were cultured on coverslips and allowed to spread out fully. Following in situ RT reactions, tubulin antibody and DRAQ5 dye were used to label the cytoplasmic and nuclear regions, respectively. DRAQ5-positive (Nucleus) and tubulin-positive (Cytoplasm) areas of the cell were independently photo-irradiated (∼100 nuclei and ∼100 cytoplasm regions with each replicate, n=4, 4, respectively) (Figure 5A). The library was prepared in accordance with the PCL-seq method as described above.

**Figure 5.**
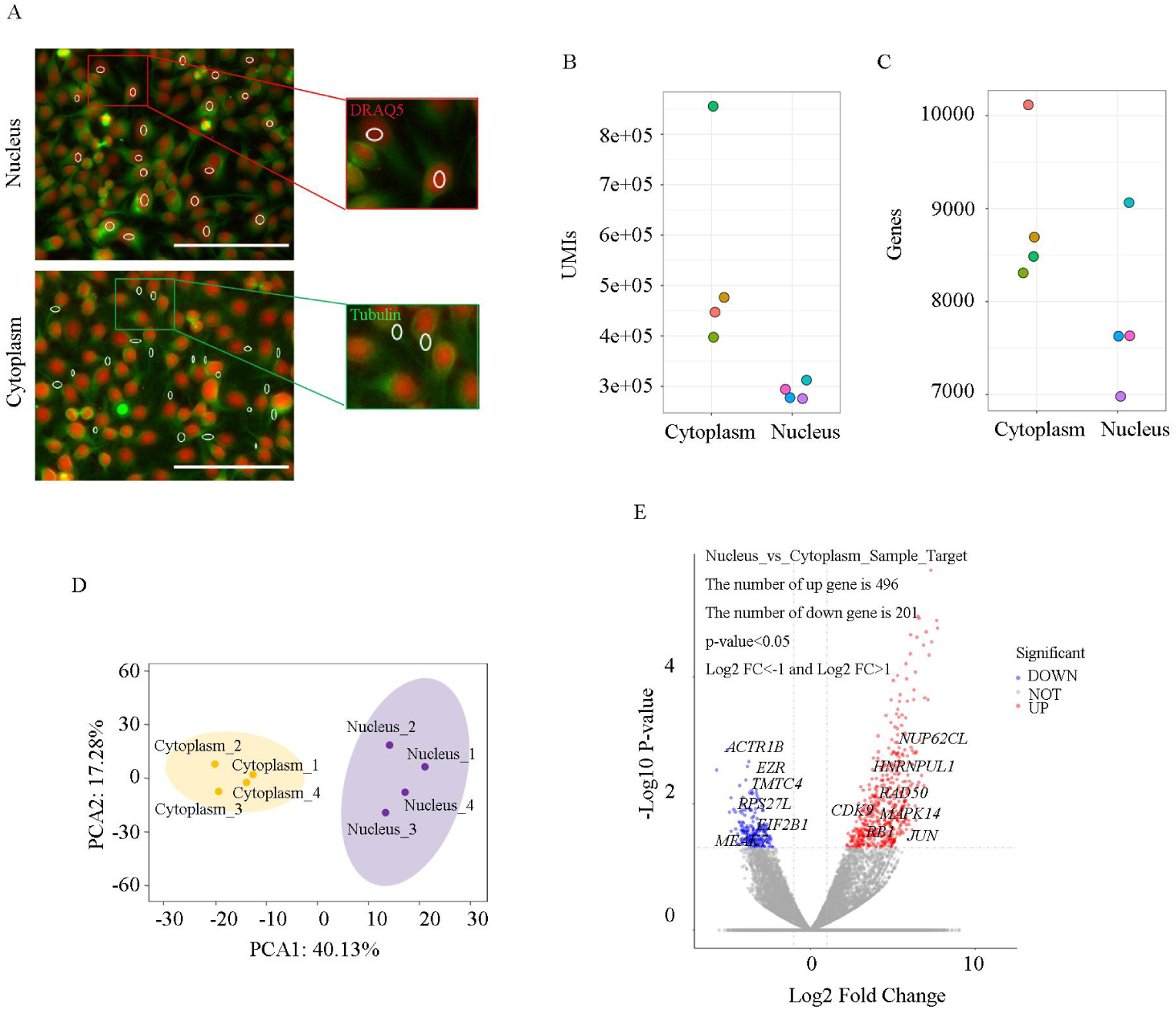
PCL-seq for subcellular structures. (A) Images of immunofluorescence staining using anti-tubulin antibody (green) for cytoplasm and DRAQ5 dye (red) for nucleus in Hela cells. Tubulin-positive and DRAQ5-positive areas were photo-irradiated. Scale bar, 200μm. (B, C) The number of detected UMIs (B) and genes (C) are shown. (D) Two-dimensional PCA for the expression profiles is shown. (E) DEGs are clustered and shown in volcano plots.

Firstly, PCL-seq exhibits excellent data output capabilities in the analysis of subcellular regions.With ∼1.7 × 10^7^ reads per sample. It produced an average of 5.4 × 10^5^ and 2.9 × 10^5^ UMIs in tubulin-positive and DRAQ5-positive regions, respectively (Figure 5B). Additionally, an average of 8,900 and 7,826 genes were detected in the two regions, respectively (Figure 5C).

Moreover, two dimensional reduction of the expression profiles suggested that DRAQ5-posi and tubulin-posi regions were well distinguished (Figure 5D). DEG analysis revealed a total of 697 genes in DRAQ5-posi vs. Tubulin-posi samples (Figure 5E). Theoretically, UV light will inevitably pass through the cytoplasm during nucleus-specific irradiation in the DRAQ5-positive specimens, so it is understandable that the DRAQ5 group has both nuclear genes and a small number of cytoplasmic genes. Despite this, we still detected the presence of *RAD50*, *NUP62CL*, *HNRNPUL1*, *MAPK14*, *CDK9*, *RB1* and *JUN* in the DRAQ5-positive group. The nuclear localization of these genes has been confirmed by the Human Protein Atlas (HPA) database, which provides tissue and cellular distribution information for all 24,000 human proteins. For example, *RAD50* is involved in the processing of double-strand break repair and DNA damage checkpoint activation^45^. *NUP62CL* is a structural component of the nuclear pore complex, involved in RNA export and protein import^46^. Simultaneously, within the tubulin-positive samples, we also detected cytoplasm-specific localization genes that have been validated in the HPA database, including *EZR*, *EIF2B1*, *RPS27L*, *TMTC4* and so on. In conclusion, these results indicate that PCL-seq can perform high-depth analyses of gene expression profiles in subcellular regions.

## Discussion

Clinical pathology tissue samples, especially those derived from complex organs such as tumors or the brain, often exhibit significant heterogeneity. This intrinsic variability makes it challenging to obtain accurate and relevant data through conventional analysis methods. Consequently, spatial transcriptomic analysis of ROI becomes essential. By focusing on specific anatomical regions or cell types, ROI analysis enables researchers to obtain highly specific target gene expression information. This targeted approach minimizes the complexities inherent in large-scale data analysis across entire samples, enhancing both the accuracy and relevance of the findings. In the context of pathological research, ROI analysis offers substantial advantages. For instance, it facilitates the identification and comparison of gene expression differences between normal and diseased regions. Such detailed comparisons can unveil disease-related molecular mechanisms that may not be apparent through broader analytical techniques. Furthermore, by pinpointing these molecular mechanisms, ROI analysis provides more precise molecular targets for personalized treatment strategies. This precision enables the development of tailored therapies that are specifically designed to address the unique molecular characteristics of an individual’s disease, thereby enhancing treatment efficacy and patient outcomes.

Here, we present PCL-seq, a method to attach sequenceable spatial adaptors to mRNA biomolecules based on the properties of PC linker, which shed after irradiation, enabling the analysis of transcriptomic profiles within ROI. We demonstrated that PCL-seq can be used for transcriptome profiling of flexible-sized ROI within both fresh frozen and FFPE tissue sections, achieving sensitivity comparable to existing methods (Supplementary data)^27,28^. Notably, PCL-seq can analyze transcription profiles of subcellular regions with high depth, which significantly advances the understanding of gene functions and regulatory mechanisms. PCL-seq utilizes a PC linker as a photo-controllable target, substantially reducing the cost of spatial transcriptomics and making the technique more accessible. The cost of primers modified with PC linker is only USD 77 (2 OD), a stark contrast to the USD 2,944 (1 OD) required for NPOM-caged primers used in PIC^27^. Additionally, the application of Light-seq is constrained by the non-commercial availability of CNVK compounds^28^. Another approach, the GeoMx DSP platform, also employs PC linker to achieve highly multiplexed spatial profiling of proteins or RNAs through iterative UV-cleavage and microcapillary collection of released PC oligonucleotides^25^. However, the DSP platform faces limitations due to its dependence on specific probes, which significantly restricts the exploration of non-human and non-murine species, as well as unknown genes. Moreover, the high cost of the GeoMx system’s experimental equipment is prohibitive for many laboratories. In summary, PCL-seq leverages the PC linker in conjunction with the Mosaic system to establish a highly reproducible and high-performance spatial transcriptomics technique, making it feasible for standard laboratory usage. This method represents a significant advancement in the field, combining cost-effectiveness with robust analytical capabilities.

In multicellular systems, spatially specific gene expression is orchestrated by chromatin conformation and epigenetic modifications^47,48^. Therefore, we anticipate that PCL-seq could extend beyond merely studying the spatial transcriptome. A novel approach, Photoselective sequencing, integrates the PC linker with ATAC-seq^49^ to facilitate spatial analysis of ROI at the chromatin level^50^. This implies that introducing the PC linker into both the Tn5 mosaic end adapter and the RT primer could enable spatial multi-omics studies of chromatin accessibility and transcriptome within the same tissue section. Moreover, PC linker can be modified into the sequence of the PA-Tn5 enzyme^51^ or antibody-oligo complex^52^, allowing for the construction of spatially resolved epigenetic and proteomic data. This advancement would provide deeper insights into the intricate regulation of gene expression and the interaction between different layers of cellular information. Such integration could pave the way for comprehensive spatially resolved multi-omics analyses, ultimately enhancing our understanding of cellular heterogeneity and tissue architecture.

## Methods

### Cell

HEK293, NIH/3T3 and Hela cells were cultured according to standard procedures in high-glucose DMEM (Gibco) supplemented with 10% FBS (Gibco), and 1% penicillin-streptomycin (Gibco) at 37℃ with 5% CO_2_. To prepare cell experiments, we seeded the cells onto a poly-L-lysine coated glass slides (Solarbio). In brief, cells were collected from culture dish by trypsin (Gibco). Cell suspension after centrifuged dripped into a slide and plated for overnight at 37℃. The slides were then fixed with 4% formaldehyde (Sigma).

### Mice

C57BL/6J mice were purchased from Jinan Pengyue Laboratory Animal Breeding Co., LTD. The study was approved by the Animal Care and Use Committee of Ocean University of China. E13.5 mouse embryos were dissected, then transferred into a cryomold previously filled with pre-chilled OCT Compound (Sakura) for embedding. The cryomold was immediately placed on dry ice until frozen. Before sectioning, the frozen tissue block was warmed to the temperature of cryotome cryostat (−20℃). Cryosections at a thickness of 10 μm were mounted on poly-L-lysine coated glass slides. The frozen slides were then fixed with 4% formaldehyde or directly kept at −80℃ if a long-time storage is needed.

Additionally, to obtain FFPE samples, E15.5 mouse embryos were fixed in 4% formaldehyde at 4°C overnight. Following fixation, the embryos were processed for paraffin embedding and sectioning Wuhan Servicebio Technology CO., LTD to provide FFPE biological samples for subsequent research.

### Functionalized slide screening for optimal irradiation conditions

First, the NH_2_-RT-PC linker-FAM primer was conjugated to the surface of CodeLink slides. Briefly, the primer was dissolved in 10 mM sodium phosphate buffer to prepare a 100 μM stock solution. The print buffer was prepared as follows: 300 mM sodium phosphate buffer pH 8.5, and 10 μM NH_2_-RT-PC linker-FAM primer. The print buffer was applied to the CodeLink slides, and the incubation area was marked. The slides were then placed in a saturated sodium chloride humid chamber and incubated overnight at room temperature in the dark. After the overnight incubation, the slides were treated with a preheated blocking solution (50 mM Ethanolamine, 0.1 M Tris pH 9) to block any remaining active groups. The blocking was carried out at 50°C for 30 minutes. Following the blocking step, the slides were briefly rinsed with deionized water. Next, 10 ml of preheated washing solution (4× SSC, 0.1% SDS) at 50°C was added to the slides, which were then washed on a shaker for 30 minutes. After a brief rinse with deionized water, the slides were dried, and the presence of green fluorescence in the reaction area under a microscope indicated successful conjugation. Subsequently, different areas of the slides were exposed to UV light at varying intensities (10%, 40%, 70%, 100% power) for different durations (10, 20, 30 seconds). After washing with PBS, the slides were observed under a fluorescence microscope, and the fluorescence intensity was recorded for each condition.

### Fluorescent labeling of ROI

Two fluorescently labeled primers (RT-PC linker-FAM and Ligation adaptor-ROX) were employed to complete CodeLink slides and tissue sections staining within the ROI. For CodeLink slides, the RT-PC linker-FAM primer can be conjugated to the surface of the slide using the aforementioned method, and subsequently exposed to UV light directly. For tissue sections, the fluorescently modified primers needs to be introduced via an RT reaction, followed by downstream processing. Briefly, tissues were fixed with 4% formaldehyde and subsequently permeabilized with 0.1% Triton X-100. A hybridization buffer containing 2× SSC (Invitrogen), 10% formamide (Solarbio), 50% Dextran sulfate (Sigma), 1% RNase inhibitor (Takara), and 1 μM RT-PC linker-FAM primer was applied to the samples for 30 minutes at room temperature. The slides were then washed three times with 2× SSC, followed by incubation with an RT mix containing 1× RT buffer (Thermo Fisher), 100 μM dNTPs (Thermo Fisher), 1% RNase inhibitor, and 25 U/μl Maxima H Minus Reverse Transcriptase (Thermo Fisher) for 90 minutes at 42°C. After washing with PBS, the target region was illuminated with 365 nm UV light at 100% power for 10 seconds. Subsequently, the slides were rinsed continuously with PBS for 3 minutes, and then 1 μM ligation adaptor-ROX primer (partially double-stranded by annealing of two single strands, linker and Ligation-ROX) was hybridized to the samples for 15 minutes at room temperature. After washing with 2× SSC, a ligation mix containing 1× T4 buffer (NEB) and 20 U/μl T4 DNA ligase (NEB) was added to the samples and incubated for 30 minutes at 37°C. Imaging was performed using a Nikon ECLIPSE Ti2 inverted microscope after washing with 65% formamide.

### PCL-seq protocol

#### Fixation and permeabilization

Fresh frozen tissue sections were fixed with 4% formaldehyde for 10 minutes at room temperature and then the cross-link was quenched by 1.25 M glycine for 5 minutes. Sections were permeabilized with 0.5% Triton X-100 for 3 minutes and then with 0.1 N HCl for 5 minutes, followed by neutralization with 1 M Tris-HCl, pH8.0, for 10 minutes at room temperature.

Typically, cells are fixed with 4% formaldehyde at room temperature for 10 minutes, followed by permeabilization with 0.1% Triton X-100 at room temperature for 20 minutes. These conditions are used for the pre-treatment of cell coverslips. Downstream processing is consistent with that of tissue sections.

#### Preparation of FFPE samples

E15.5 mouse embryo FFPE sections (10 μm) were deparaffinized by immersion in the following reagent: Xylene (VWR) 15 minutes twice, EtOH 99% (VWR) 2 minutes twice, EtOH 96% (VWR) 2 minutes twice, EtOH 70% (VWR) 2 minutes twice, H_2_O 5 minutes. Then, a collagenase mix containing 1% BSA, 50U/μl collagenase I was added and incubated for 20 minutes at 37℃. Once the incubation was complete, the collagenase mix was washed by 0.1× SSC buffer. Subsequently, TE buffer pH8.0 was added, then the slides were incubated for 1 hour at 70℃. After the incubation, the slides left to equilibrate at room temperature for 5 minutes. Meanwhile, 0.1% pepsin solution dissolved in 0.1 M HCl was equilibrated to 37℃. After the incubation and wash with 0.1× SSC buffer, permeabilization of the tissue was carried out by adding pepsin solution and incubating at 37℃ for 30 minutes. After this step, 0.1× SSC buffer was added to wash the pepsin solution. The next steps are the same as formaldehyde-fixed tissue sections.

#### Reverse transcription

RT-PC linker primers were hybridized to the tissue section by applying a hybridization buffer solution consisting of 2× SSC, 10% formamide, 50% Dextran Sulfate, and 1% RNase inhibitor onto the slide. The hybridization was incubated for 30 minutes at room temperature. Subsequently, the section was rinsed three times with 2× SSC and once with 1× RT buffer. Following these rinses, the section was subjected to a reverse transcription mix containing 1× RT buffer, 100 μM dNTPs, 1% RNase inhibitor, and 25 U/μL Maxima H Minus Reverse Transcriptase for 90 minutes at 42°C.

#### Immunofluorescence staining (optional)

For tissue sections, Alexa Fluor® 647 anti-PAX6 antibody (Abcam) was used to visualize the eye structure of mouse embryos. To prevent non-specific protein-protein interactions, the specimens were blocked with 1% BSA for 1 hour at room temperature, followed by three washes with PBST. Subsequently, the tissue sections were incubated with the anti-PAX6 antibody for 2 hours at room temperature. After incubation, the sections were rinsed three times with PBST to complete the fluorescent staining. Based on the staining results, ROI can then be selected.

For HeLa cells, immunofluorescence staining is performed using an anti-tubulin antibody to visualize the cytoplasmic region. Subsequently, staining with DRAQ5^TM^ (Invitrogen) is used to indicate the nuclear location. Briefly, post-RT reaction cells are blocked with 1% BSA for 1 hour at room temperature. After three washes with PBST, the cells are incubated with the Alexa Fluor® 488 anti-tubulin antibody (Abcam) for 2 hours at room temperature to stain the cytoplasm. Following another PBST wash, the cells are incubated with 10 μM DRAQ5^TM^ dye for 30 minutes at room temperature. The cells are then washed with PBST to prepare for subsequent reactions.

#### Blocking reaction

To prevent large tissue areas from being ligated with the adaptor due to background shedding of the PC Linker during primer synthesis, blocking primers were employed to obstruct RT primers lacking the PC Linker. Briefly, 10 μM linker strands complementary to the RT primer and 10 μM block primers (non-amplification arm strands) were hybridized to the section for 15 minutes at room temperature. Following these single-strand hybridizations, the slides were washed with 2× SSC and subsequently with 1× T4 buffer. A ligation mix containing 1× T4 buffer and 20 U/μL T4 DNA ligase was then applied to the section and incubated for 30 minutes at 37°C. Finally, the slides were washed with 65% formamide to remove single strands complementary to the RT primers, allowing for subsequent specific primer hybridization in the irradiated area.

#### Photo-cleavable of PC Linker

Photo-cleavable of PC Linker was performed with a customized Nikon ECLIPSE Ti2 inverted microscope fitted with a Mosaic Digital Mirror Devices (DMD) from Andor. The DMD comprises an array of individually addressable micro-mirrors that can be switched “on and off”. In the imaging software, our defined ROI is converted into a mask, which is reflected in the micro-mirrors. This patterned light is then directed through the microscopy and objective onto the sample. In our experiments, a slide attached tissue is facing the objective, so that UV light can shine directly on the tissue without passing through the glass to avoid light deflection. Subsequently, the desired ROI were illuminated using 100% power for 10 seconds to accomplish photo-cleavable and the broken PC Linker were thoroughly cleaned with PBS.

#### Ligation reaction

After rinsing with PBS continuously for 3 minutes, 1 μM linker strands and 1 μM ligation strands were hybridized to the section for 15 minutes at room temperature. Subsequently, the slides were washed with 2×SSC and then with 1× T4 buffer. A ligation mix, consisting of a final concentration of 1× T4 buffer and 20 U/μL T4 DNA ligase, was added to the section and incubated for 30 minutes at 37°C. Following incubation, the slides were washed with 65% formamide to prevent nonspecific hybridization.

#### cDNA collection and purification

An incomplete reverse-crosslinking solution, containing 50 mM Tris-HCl, 1 mM EDTA, 0.2 M NaCl, and 1 mg/mL proteinase K, was added to the tissue section for 2 minutes at 55°C. The mixture was then carefully transferred to an Eppendorf tube. The glass slide was washed three additional times with the same buffer, and the wash solutions were also carefully transferred to the same tube. After adding SDS to a final concentration of 1% to the tissue lysate, the mixture was incubated overnight at 55°C. The reverse-crosslinked product was then purified using Ampure XP beads (Beckman) at a 0.6× ratio. The mRNA-cDNA complex was collected from the beads and further purified using Dynabeads MyOne Streptavidin C1 beads (Thermo Fisher). The streptavidin beads were first washed three times with binding buffer containing 10 mM Tris-HCl pH 7.5, 1 mM EDTA, and 1 M NaCl, and then resuspended in the same buffer. The product purified by the Ampure XP beads was transferred to a tube containing the washed streptavidin beads. The mixture was vigorously mixed at 1600 rpm for 30 minutes at room temperature. Finally, the beads were thoroughly washed with wash buffer 1 (1× SSC, 0.1% SDS) once for 15 minutes at room temperature and wash buffer 2 (0.1× SSC, 0.1% SDS) three times for 10 minutes at 65°C.

#### Template switching and PCR amplification

The cDNAs bound to the beads were resuspended in the template switching solution, which contained 1× RT buffer, 1 mM dNTPs, 1 μM TSO primer, 1% RNase inhibitor, and 2 U/μL Maxima H Minus Reverse Transcriptase. The reaction was performed at 42°C for 90 minutes. Subsequently, a mixture containing KAPA HIFI HotStart Master Mix (Kapa Biosystems) with 1× KAPA buffer, 0.3 M dNTPs, 0.2 mM MgCl2, 0.4 μM ISPCR primer, and 1 U/μL HotStart HIFI enzyme was added to the template switching product. The PCR reaction was carried out with the following conditions: an initial incubation at 98°C for 3 minutes, followed by 23 cycles of 98°C for 20 seconds, 67°C for 10 seconds, and 72°C for 6 minutes, concluding with a final extension at 72°C for 5 minutes. The PCR product was then purified using Ampure XP beads at a 0.6× ratio and eluted into 10 μL of 10 mM Tris-HCl, pH 8.0 buffer. The quantity of cDNAs was measured using the Qubit fluorometer (Invitrogen).

#### Library preparation

To construct the sequencing library, the purified cDNAs were diluted to a concentration of 1 ng/μL and subjected to tagmentation with Tn5A transposase for 10 minutes at 55°C. The tagmentation reaction was quenched by the addition of 0.2% SDS, and the product was purified using Ampure XP beads at a 1× ratio. The cDNAs were then amplified using KAPA HIFI Master Mix (Kapa Biosystems) comprising 1× KAPA buffer, 0.3 M dNTPs, 0.2 mM MgCl2, 0.2 μM P7 primer, 0.2 μM S5 primer, and 1 U/μL HIFI enzyme. The PCR amplification conditions were as follows: an initial incubation at 72°C for 5 minutes, followed by 15 cycles of 95°C for 20 seconds, 55°C for 30 seconds, and 72°C for 30 seconds, and a final extension at 72°C for 5 minutes. The PCR product was further purified using Ampure XP beads. Sequencing was performed on the Illumina NovaSeq platform by Novogene.

### Data analysis

#### Library screening and analysis workflow

The CellCosmo 1.0.11 software’s rna barcode module was utilized to filter and obtain reads conforming to the library structure. Subsequently, these reads were subjected to standard analysis using the Drop-seq software. The overall workflow includes the following main steps:

1. Barcode splitting (TagBamWithReadSequenceExtended module);
2. Filtering low-quality sequences (PolyATrimmer module);
3. Reference genome alignment (using STAR software);
4. Quantification of reads to exon+intron (TagReadWithGeneFunction module);
5. Barcode correction based on UMI counts (DetectBeadSynthesisErrors module);
6. Generation of barcode-gene expression matrix (DigitalExpression module).

#### Proportion analysis of human and mouse genic reads

The upstream processing employed the CellCosmo rna barcode module and Drop-seq’s standard workflow, with a modification in the reference genome. Specifically, the workflow incorporated a combined reference genome of human and mouse, with gene names and chromosome identifiers prefixed by “hg38_” for human and “mm10_” for mouse. Before the step of generating the barcode-gene expression matrix, the FilterBam module was used to split the BAM file into separate human and mouse files based on the chromosome prefix “hg38” and “mm10”. Subsequently, the DigitalExpression module was used to generate two single-cell gene expression matrices: one for human and one for mouse. The proportion analysis of genic reads was then conducted separately for human and mouse.

#### Differential gene expression analysis

Differentially expressed genes (DEGs) were identified using DESeq2 for pairwise comparison analysis. For this purpose, Seurat v3.1.2’s FindMarkers function was utilized with default parameters, employing the Wilcoxon likelihood-ratio test. Genes were selected as differentially expressed if they were expressed in more than 10% of cells within a cluster and had an average log2 fold-change greater than 0.25. Analytical tools and visualization methods:

Differential gene expression analysis: Conducted using the DESeq2 package in R. Volcano plot: Created with the ggplot2 package.

Principal component analysis (PCA): Performed using the Gmodels and FactoMineR packages.

Heatmap: Generated using the Pheatmap2 package.

## Data availability statement

All raw and processed sequencing data generated in this study have been submitted to the NCBI Gene Expression Omnibus (GEO; https://www.ncbi.nlm.nih.gov/geo/) under accession number GSExxxxxx. Upon acceptance of the manuscript, we pledge to release the data stored in the database to the public domain.

## Acknowledgment

This work is supported by the Funding Project of National Key Research and Development Program of China (2018YFD0900604) and Natural Science Foundation of China (41676119 and 41476120).

---

**Supplementary Figure 1.**
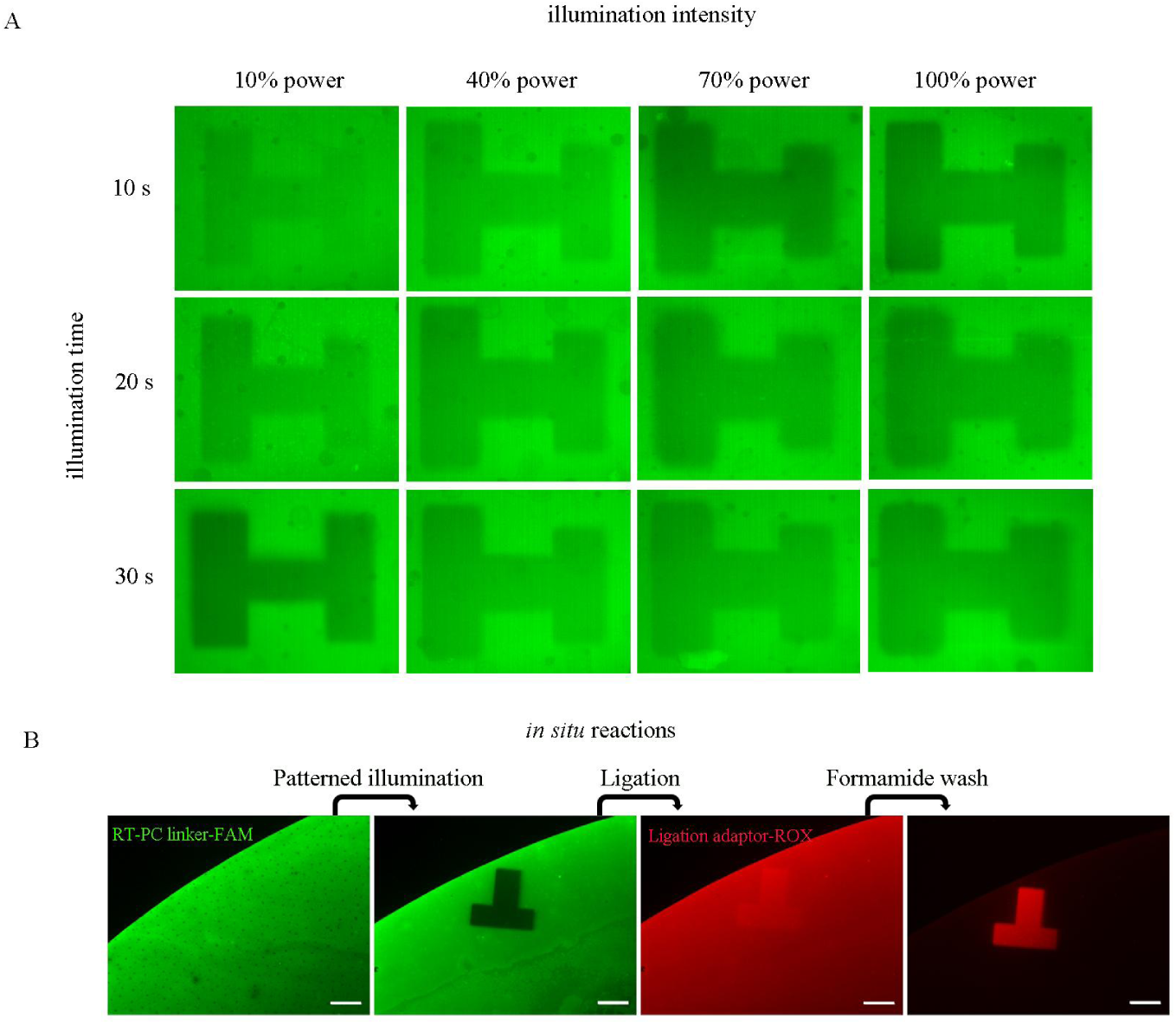
Optimization of specific ROI fluorescent labeling in PCL-seq. (A) Different light intensities and illumination times were tested to determine the best exposure condition. The area marked with ‘H’ denotes the regions that were subjected to UV irradiation. (B) A fluorescence pattern is observed on CodeLink slides post-UV illumination, ligation, and formamide rinsing. The T-shaped area is the photo-activated region. Scale bar, 100 μm.

**Supplementary Figure 2.**
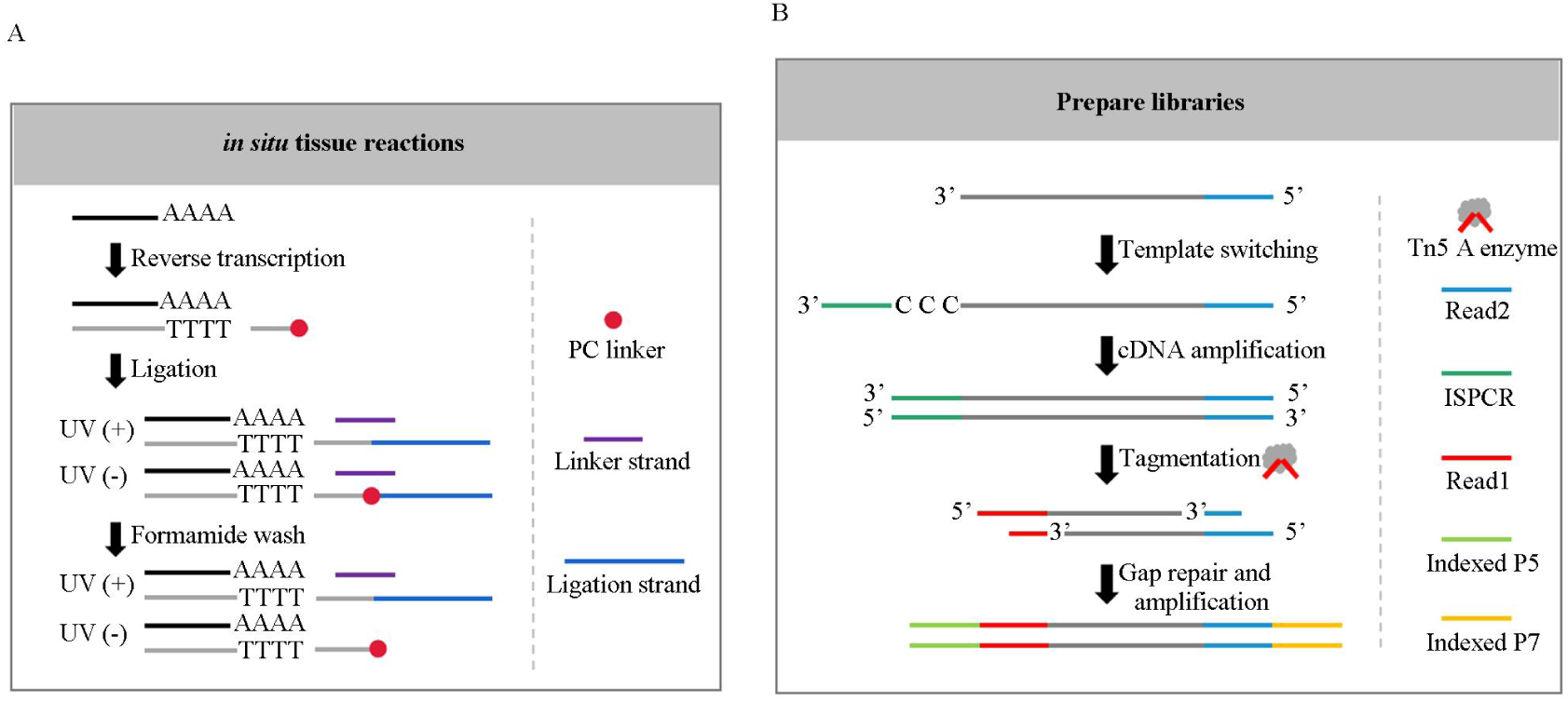
Schematic diagram of in situ reactions and library preparation for PCL-seq. (A) In situ tissue reaction: RT primers modified with a PC linker are used to capture mRNA and facilitate reverse transcription within the tissue. The ROI is exposed to UV light, which cleaves the PC linker at the 5’-terminal of the RT primer, permitting the mRNA-cDNA complex to ligate with a sequencing adapter. In regions illuminated by UV light, this ligation is stable and resistant to formamide removal. In contrast, in non-illuminated areas, the PC linker prevents ligation of the sequencing adapter. The linker strand in these areas remains hybridized to the complementary sequence on the RT primer and can be removed by formamide treatment. (B) Library construction: Following tissue digestion and purification, a second reverse transcription reaction is performed. This involves amplification of both ends of the cDNA using ISPCR primers. The sample is then treated with Tn5 transposase, which is equipped only with Mosaic end adaptor A. This enzyme not only fragments the cDNA but also initiates the insertion of sequencing adapters. A subsequent gap-fill step ensures the generation of complete double-stranded DNA fragments. The library is then amplified from these ligated adapters, preparing it for next-generation sequencing.

**Supplementary Figure 3.**
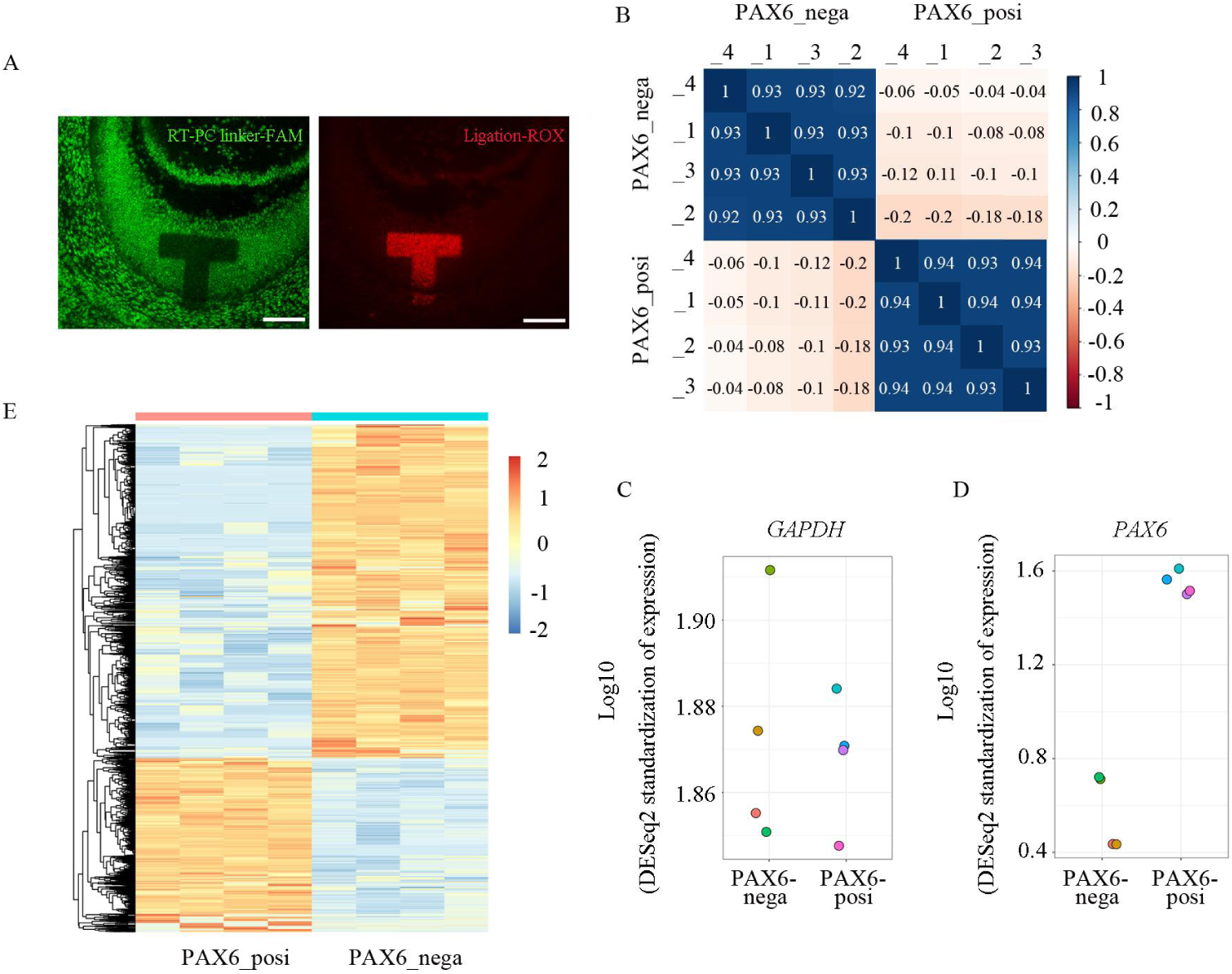
PCL-seq for fresh frozen mouse embryo sections. (A) A mouse embryo section was stained with RT-PC linker-FAM primers via an in-situ RT reaction and subsequently imaged after photo-cleavage, ligation, and formamide rinsing. The T-shaped area indicates the photo-activated region. Scale bar, 100 μm. (B)The correction of different specimens. (C, D) The expression of *GAPDH* (C) and *PAX6* (D) genes were examined. (E) DEGs are clustered and shown in heat maps.

**Supplementary Figure 4.**
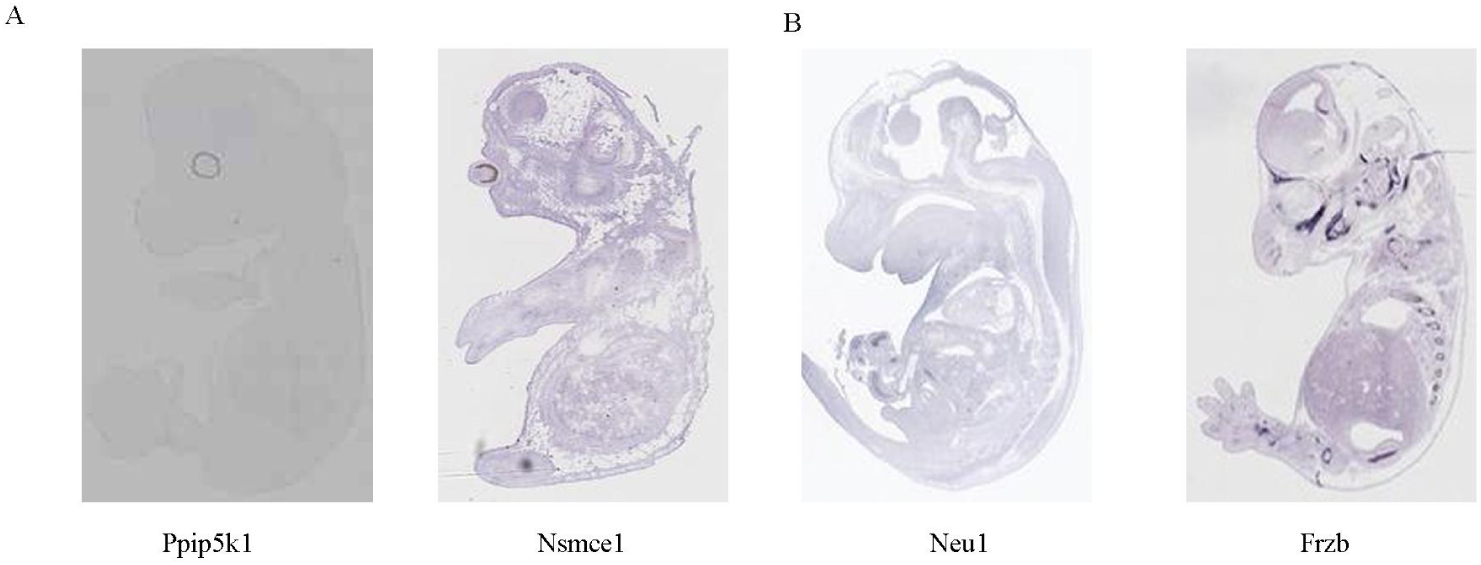
ISH experiments by EMAGE with differential expression genes. ISH experiments are demonstrated that the expression levels of these genes are indeed different in the eye (A), limb and skeleton (B) part of mouse embryo.

**Supplementary Table 1:**
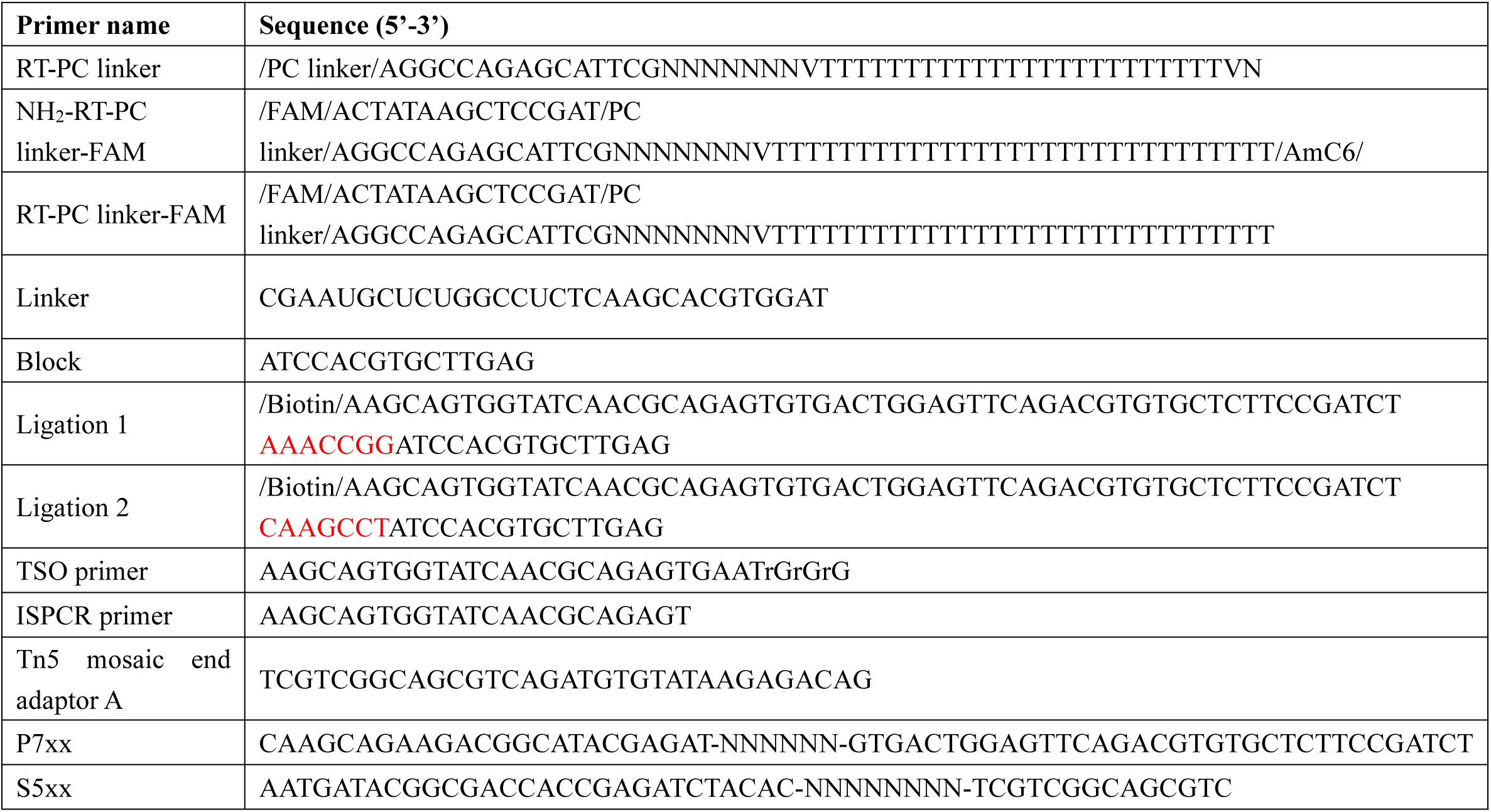
Primer names and sequences of PCL-seq. The purification method for primers shown in the table is HPLC.

## References

1 Wang, F., et al. Single-cell and spatial transcriptome analysis reveals the cellular heterogeneity of liver metastatic colorectal cancer. Sci Adv 9, 16 (2023).

2 Hwang, W. L., et al. Single-nucleus and spatial transcriptome profiling of pancreatic cancer identifies multicellular dynamics associated with neoadjuvant treatment. Nat Genet 54, 1178–1191 (2022).

3 Hildebrandt, F., et al. Spatial Transcriptomics to define transcriptional patterns of zonation and structural components in the mouse liver. Nat Commun 12, 021–27354 (2021).

4 Tavares-Ferreira, D., et al. Spatial transcriptomics of dorsal root ganglia identifies molecular signatures of human nociceptors. Sci Transl Med 14, 16 (2022).

5 Chen, A., et al. Spatiotemporal transcriptomic atlas of mouse organogenesis using DNA nanoball-patterned arrays. Cell 185, 1777–1792 (2022).

6 Delile, J., et al. Single cell transcriptomics reveals spatial and temporal dynamics of gene expression in the developing mouse spinal cord. Development 146, 173807 (2019).

7 Liu, Y., et al. Integration analysis of single-cell and spatial transcriptomics reveal the cellular heterogeneity landscape in glioblastoma and establish a polygenic risk model. Front Oncol 13 (2023).

8 Shiau, C., et al. Therapy-associated remodeling of pancreatic cancer revealed by single-cell spatial transcriptomics and optimal transport analysis. (bioRxiv. 2023 Jun 29:2023.06.28.546848. doi: 10.1101/2023.06.28.546848. Preprint.).

9 Moncada, R., et al. Integrating microarray-based spatial transcriptomics and single-cell RNA-seq reveals tissue architecture in pancreatic ductal adenocarcinomas. Nat Biotechnol 38, 333–342 (2020).

10 Melo Ferreira, R., et al. Integration of spatial and single-cell transcriptomics localizes epithelial cell-immune cross-talk in kidney injury. JCI Insight 6, 147703 (2021).

11 van den Brink, S. C., et al. Single-cell and spatial transcriptomics reveal somitogenesis in gastruloids. Nature 582, 405–409 (2020).

12 Rahman, S. & Zenklusen, D. Single-molecule resolution fluorescent in situ hybridization (smFISH) in the yeast S. cerevisiae. Methods Mol Biol, 526–522_523 (2013).

13 Shah, S., Lubeck, E., Zhou, W. & Cai, L. In Situ Transcription Profiling of Single Cells Reveals Spatial Organization of Cells in the Mouse Hippocampus. Neuron 92, 342–357 (2016).

14 Eng, C. L., et al. Transcriptome-scale super-resolved imaging in tissues by RNA seqFISH. Nature 568, 235–239 (2019).

15 Chen, K. H., Boettiger, A. N., Moffitt, J. R., Wang, S. & Zhuang, X. RNA imaging. Spatially resolved, highly multiplexed RNA profiling in single cells. Science 348, 9 (2015).

16 Ståhl, P. L., et al. Visualization and analysis of gene expression in tissue sections by spatial transcriptomics. Science 353, 78–82 (2016).

17 Rodriques, S. G., et al. Slide-seq: A scalable technology for measuring genome-wide expression at high spatial resolution. Science 363, 1463–1467 (2019).

18 Stickels, R. R., et al. Highly sensitive spatial transcriptomics at near-cellular resolution with Slide-seqV2. Nat Biotechnol 39, 313–319 (2021).

19 Vickovic, S., et al. High-definition spatial transcriptomics for in situ tissue profiling. Nat Methods 16, 987–990 (2019).

20 Srivatsan, S. R., et al. Embryo-scale, single-cell spatial transcriptomics. Science 373, 111–117 (2021).

21 Moses, L. & Pachter, L. Museum of spatial transcriptomics. Nat Methods 19, 534–546 (2022).

22 Genshaft, A. S., et al. Live cell tagging tracking and isolation for spatial transcriptomics using photoactivatable cell dyes. Nat Commun 12, 021–25279 (2021).

23 Nichterwitz, S., et al. Laser capture microscopy coupled with Smart-seq2 for precise spatial transcriptomic profiling. Nat Commun 7 (2016).

24 Chen, J., et al. Spatial transcriptomic analysis of cryosectioned tissue samples with Geo-seq. Nat Protoc 12, 566–580 (2017).

25 Merritt, C. R., et al. Multiplex digital spatial profiling of proteins and RNA in fixed tissue. (Nat Biotechnol. 2020 May;38(5):586–599. doi: 10.1038/s41587-020-0472-9. Epub 2020 May 11.).

26 Hu, K. H., et al. ZipSeq: barcoding for real-time mapping of single cell transcriptomes. Nature Methods 17, 833-+, doi:10.1038/s41592-020-0880-2 (2020).

27 Honda, M., et al. High-depth spatial transcriptome analysis by photo-isolation chemistry. Nat Commun 12, 021–24691 (2021).

28 Kishi, J. Y., et al. Light-Seq: light-directed in situ barcoding of biomolecules in fixed cells and tissues for spatially indexed sequencing. Nat Methods 19, 1393–1402 (2022).

29 Olejnik, J., Krzymanska-Olejnik, E. & Rothschild, K. J. Photocleavable aminotag phosphoramidites for 5’-termini DNA/RNA labeling. Nucleic Acids Res 26, 3572–3576 (1998).

30 Wu, P. & Grainger, D. W. Toward immobilized antibody microarray optimization: print buffer and storage condition comparisons on performance. Biomed Sci Instrum 40, 243–248 (2004).

31 Heavner, W. & Pevny, L. Eye development and retinogenesis. Cold Spring Harb Perspect Biol 4 (2012).

32 Smith, A. N., Miller, L. A., Radice, G., Ashery-Padan, R. & Lang, R. A. Stage-dependent modes of Pax6-Sox2 epistasis regulate lens development and eye morphogenesis. Development 136, 2977–2985 (2009).

33 Love, M. I., Huber, W. & Anders, S. Moderated estimation of fold change and dispersion for RNA-seq data with DESeq2. Genome Biol 15, 014–0550 (2014).

34 Liu, Y., et al. High-Spatial-Resolution Multi-Omics Sequencing via Deterministic Barcoding in Tissue. Cell 183, 1665–1681 (2020).

35 Karia, B., Martinez, J. A. & Bishop, A. J. Induction of homologous recombination following in utero exposure to DNA-damaging agents. DNA Repair 12, 912–921 (2013).

36 Wang, N. K., et al. Transplantation of reprogrammed embryonic stem cells improves visual function in a mouse model for retinitis pigmentosa. Transplantation 89, 911–919 (2010).

37 Shears, S. B., Baughman, B. M., Gu, C., Nair, V. S. & Wang, H. The significance of the 1-kinase/1-phosphatase activities of the PPIP5K family. Adv Biol Regul 63, 98–106 (2017).

38 Gong, M., et al. A transcriptomic analysis of Nsmce1 overexpression in mouse hippocampal neuronal cell by RNA sequencing. Funct Integr Genomics 20, 459–470 (2020).

39 Fassunke, J., et al. Utility of different massive parallel sequencing platforms for mutation profiling in clinical samples and identification of pitfalls using FFPE tissue. Int J Mol Med 36, 1233–1243 (2015).

40 Hoffman, E. A., Frey, B. L., Smith, L. M. & Auble, D. T. Formaldehyde crosslinking: a tool for the study of chromatin complexes. J Biol Chem 290, 26404–26411 (2015).

41 Gracia Villacampa, E., et al. Genome-wide spatial expression profiling in formalin-fixed tissues. Cell Genom 1, 8 (2021).

42 Bai, Z. et al. Spatially Exploring RNA Biology in Archival Formalin-Fixed Paraffin-Embedded Tissues. (bioRxiv [Preprint]. 2024 Feb 8:2024.02.06.579143. doi: 10.1101/2024.02.06.579143.).

43 Turner-Bridger, B., et al. Single-molecule analysis of endogenous β-actin mRNA trafficking reveals a mechanism for compartmentalized mRNA localization in axons. Proc Natl Acad Sci U S A 115, E9697–E9706 (2018).

44 Das, S., Vera, M., Gandin, V., Singer, R. H. & Tutucci, E. Intracellular mRNA transport and localized translation. Nat Rev Mol Cell Biol 22, 483–504 (2021).

45 Li, Y., et al. Rad50 promotes ovarian cancer progression through NF-κB activation. J Cell Mol Med 25, 10961–10972 (2021).

46 Yang, Z., et al. Comprehensive analysis of resistance mechanisms to EGFR-TKIs and establishment and validation of prognostic model. J Cancer Res Clin Oncol 149, 13773–13792 (2023).

47 Peixoto, P., Cartron, P. F., Serandour, A. A. & Hervouet, E. From 1957 to Nowadays: A Brief History of Epigenetics. Int J Mol Sci 21 (2020).

48 Holoch, D. & Moazed, D. RNA-mediated epigenetic regulation of gene expression. Nat Rev Genet 16, 71–84 (2015).

49 Buenrostro, J. D., Giresi, P. G., Zaba, L. C., Chang, H. Y. & Greenleaf, W. J. Transposition of native chromatin for fast and sensitive epigenomic profiling of open chromatin, DNA-binding proteins and nucleosome position. Nat Methods 10, 1213–1218 (2013).

50 Mangiameli, S. M., et al. Photoselective sequencing: microscopically guided genomic measurements with subcellular resolution. Nat Methods 20, 686–694 (2023).

51 Kaya-Okur, H. S. et al. CUT&Tag for efficient epigenomic profiling of small samples and single cells. Nat Commun 10, 019–09982 (2019).

52 Stoeckius, M., et al. Simultaneous epitope and transcriptome measurement in single cells. Nat Methods 14, 865–868 (2017).

